# Coping with darkness: The adaptive response of marine picocyanobacteria to repeated light energy deprivation

**DOI:** 10.1101/2020.10.15.341503

**Authors:** Allison Coe, Steven J. Biller, Elaina Thomas, Konstantinos Boulias, Christina Bliem, Aldo Arellano, Keven Dooley, Anna N. Rasmussen, Kristen LeGault, Tyler J. O’Keefe, Eric L. Greer, Sallie W. Chisholm

## Abstract

The picocyanobacteria *Prochlorococcus* and *Synechococcus* are found throughout the ocean’s euphotic zone, where the daily light:dark cycle drives their physiology. Periodic deep mixing events can, however, move cells below this zone, depriving them of light for extended periods of time. Here we demonstrate that *Prochlorococcus* and *Synechococcus* can adapt to tolerate repeated periods of light energy deprivation. Cyanobacterial cultures kept in the dark for 3 days and then returned to the light initially required 18-26 days to resume growth, but after multiple rounds of dark exposure the strains began to regrow after only 1-2 days. This dark-tolerant phenotype was stable and heritable; cultures retained the trait across at least 18-21 generations even when grown in a standard 13:11 light:dark cycle. We found no genetic differences between the dark-tolerant and parental strains of *Prochlorococcus* NATL2A, indicating that an epigenetic change is responsible for the adaptation. To begin to explore this possibility, we asked whether DNA methylation – an epigenetic mechanism in bacteria – occurs in *Prochlorococcus*. LC-MS/MS analysis showed that while DNA methylations, including 6mA and 5mC, are found in some other *Prochlorococcus* strains, no methylations were detected in either the parental or dark-tolerant strain used in our experiments –i.e. the NATL2A strain. These findings suggest that *Prochlorococcus* utilizes a yet-to-be-determined epigenetic mechanism to adapt to the stress of extended light energy deprivation.

## Introduction

*Prochlorococcus* are small (<1μm) non-motile cyanobacterial cells that are broadly distributed throughout the mid-latitude oceans, with a global population of ~3×10^27^ cells (Flombaum et al. 2013). Its close relative, the marine *Synechococcus*, number ~7 × 10^26^ and can also be found throughout the global oceans (Flombaum et al. 2013). *Prochlorococcus* comprises a number of genetically and physiologically distinct ecotypes, which can be broadly classified as made up of High-Light (HL) and Low-Light (LL)-adapted cells (Moore and Chisholm 1999; West and Scanlan 1999; Rocap et al. 2002). HL and LL cells exhibit distinct distributions throughout the euphotic zone, and across global spatial scales (Zinser et al. 2006; Johnson et al. 2006; Malmstrom et al. 2010), with the HL-adapted cells generally found in highest abundance near the surface, and LL-adapted cells found deeper in the euphotic zone.

Vertical mixing through convection, internal waves, mesoscale eddies, and turbulent mixing can periodically displace these photosynthetic cells below the euphotic zone, depriving them of light energy for extended periods (Denman and Gargett 1983; Falkowski et al. 1991; Thorpe 2004). Multiple studies have documented the presence of *Prochlorococcus* (DeLong et al. 2006; Jiao et al. 2014; Shibl et al. 2014) and *Synechococcus* (Sohrin et al. 2011; Miller et al. 2017; Callieri et al. 2019) at depths of 300 to 2000m – well below the euphotic zone. Observations of *Prochlorococcus* rRNA from deep samples in the western Pacific (Jiao et al. 2014), active transcription in *Prochlorococcus* and *Synechococcus* at 440m in the Gulf of Aqaba (Miller et al. 2017), and the isolation of viable *Synechococcus* cells from 750m in the anoxic Black Sea (Callieri et al. 2019) all indicate that these cyanobacteria may remain viable even in aphotic waters. The question then becomes: How long can they survive in the dark, and through what mechanisms?

Survival in darkness likely depends on a number of factors, including the length of light deprivation, ability to form resting states (Smayda and Mitchell-Innes 1974), reduce metabolic rates (Dehning and Tilzer 1989; Walter et al. 2017), draw on energy stores (Walter et al. 2017) or switch to reduced carbon energy sources (White 1974). *Prochlorococcus* and *Synechococcus* are indeed able to utilize organic carbon sources for some of their carbon and energy needs (Eiler 2006; Yelton et al. 2016; Muñoz-Marín et al. 2020) and we have shown that access to organic carbon – either supplied directly from the media or by co-cultured heterotrophs – can prolong *Prochlorococcus’* survival in extended darkness (Coe et al. 2016; Biller et al. 2018). When grown in strictly inorganic media, for example, pure cultures of *Prochlorococcus* are unable to survive more than 1.5 days of extended darkness; when co-cultured with a marine heterotroph such as *Alteromonas*, however, it can survive up to 11 days (Coe et al. 2016; Biller et al. 2018). The mechanisms through which the ‘helper-bacterium’(*sensu* Morris et al. 2008) aids dark-survival appear to involve complex cross-feeding interactions via organic carbon (Coe et al. 2016; Biller et al. 2018), as well as reduction of oxidative stress in the local environment (Morris et al. 2011, 2012; Ma et al. 2017). In contrast to *Prochlorococcus*, two strains of marine *Synechococcus*, WH7803 and WH8102, can survive at least 3 days in the dark without ‘helper-bacteria’ or the addition of organic carbon (Coe et al. 2016). While the mechanisms responsible for the differential tolerance of extended darkness is not clear, it is noteworthy that some *Synechococcus* lineages appear to endure very long periods of darkness in the field, including sunless winter months in the Arctic (Cottrell and Kirchman 2009), during deep mixing events in the Adriatic Sea (Vilibić and Šantić 2008), and for at least 30 days at 300m in Suruga Bay (Sohrin et al. 2011).

Cultures used in our dark-survival experiments to date (Coe et al. 2016) have been maintained for decades on either a diel light:dark cycle, or under continuous light. That is, over the past ~30 years or so the cell populations have not experienced repeated extended light deprivation that might occur in the wild. This made us wonder, what happens to *Prochlorococcus* cells that are repeatedly removed from their expected light:dark cycle? To examine this, we exposed HL and LL *Prochlorococcus*, as well as strains of *Synechococcus*, to repeated periods of extended darkness. We measured how long it took for the culture to resume growth after each extended dark exposure, as well as whether these multiple rounds of exposure enhanced the population’s ability to survive longer periods of darkness.

## Methods

### Culturing

All *Prochlorococcus* and *Synechococcus* cells were grown in 0.2 *μ*m filtered sterile Sargasso Sea water amended with Pro99 nutrients prepared as previously described (Moore et al. 2007). Triplicate cultures starting at a concentration of 5 × 10^6^ cells mL^−1^ to 1 × 10^7^ cells mL^−1^, were grown in a 13:11 light:dark cycle incubator with simulated dawn and dusk (Zinser et al. 2009) at 24 °C. This simulation creates gradual light transitions at sunrise by ramping light slowly up to mid‐day, remaining at peak light for 4 h, and then decreasing light to sunset over the course of 13 h. This gradual increase was important for reducing light shock on the cultures transitioning from extended darkness back into 13:11 light:dark conditions. Near optimal peak light levels for maximizing growth rate were used for all *Prochlorococcus* and *Synechococcus* strains involved and included the following combinations: MED4 (80 ± 1*μ*mol photons m^−2^ s^−1^), MIT9312 (80 ± 1 *μ*mol photons m^−2^ s^−1^), MIT9202 (72 ± 1 *μ*mol photons m^−2^ s^−1^), MIT9215 (72 ± 1 *μ*mol photons m^−2^ s^−1^), AS9601 (72 ± 1 *μ*mol photons m^−2^ s^−1^), NATL2A (37 ± 1 *μ*mol photons m^−2^ s^−1^), MIT9313 (29 ± 1 *μ*mol photons m^−2^ s^−1^), and WH8102 (76 ± 1 *μ*mol photons m^−2^ s^−1^). Growth rates were calculated by exponential regression from the log-linear portion of the growth curve. To compare changes in growth rates between parental and dark-trained lines, 3-7 growth curves were averaged and two-tailed homoscedastic *t*-tests were conducted using Microsoft Excel to determine significance.

To subject cells to extended darkness, we placed exponentially growing cultures into a 24 °C dark incubator at the end of the 13:11 light:dark cycle for varying durations. Including the 11 h of their last “natural” light:dark cycle, these cultures were in the dark for a total of 83 or 251 hrs for *Prochlorococcus* and 83, 155, 179, and 251 hrs for *Synechococcus*, which amounts to an additional 3, 6, 7, or 10 days of extended darkness, respectively. The cultures were then shifted back into the light:dark incubator at “sunrise” to reduce light shock effects, and recovery was monitored via bulk chlorophyll fluorescence measurements (10AU model, Turner Designs, Sunnyvale, California) and flow cytometry (see below). All dark sampling and measurements were done in green light using layered neutral density filters #736 and 740 (Lee Filters, Burbank, California) over a white light source, which causes minimal gene expression change in *Prochlorococcus* (Steglich et al. 2006). *Synechococcus* cultures were axenic and were tested for purity using three broths ProAC, ProMM, and MPTB (Saito et al. 2002; Morris et al. 2008; Berube et al. 2015), as well as by flow cytometry.

*Alteromonas macleodii* MIT1002 (Biller et al. 2015) was maintained on ProMM medium (Berube et al. 2015), but was spun down (10,000 ×*g* for 15 minutes) and washed twice with Pro99 medium, prior to addition into *Prochlorococcus* cultures to reduce carryover of organic carbon from ProMM. *Alteromonas macleodii* concentrations ranging between 5 × 10^5^ cells mL^−1^ to 1 × 10^6^ cells mL^−1^ were added at the onset of the experiment.

Cultures of *Alteromonas* and *Prochlorococcus* from dark-trained and parental co-cultures (*i.e.* cells that were not exposed to repeated light energy deprivation), were rendered axenic by using a serial dilution-to-extinction method previously published (Berube et al. 2015). Purity of the cultures was confirmed using all three purity broths mentioned above and by flow cytometry.

### Flow cytometry

*Prochlorococcus* cell abundance measurements by flow cytometry were prepared and processed as previously described (Zinser et al. 2006; Malmstrom et al. 2010). Samples were run on an Influx flow cytometer (Becton Dickinson, Franklin Lakes, New Jersey, U.S.A.) and excited with a blue 488 nm laser and analyzed for chlorophyll fluorescence (692/40 nm) and size (forward scatter). All samples included 2 μm diameter Fluoresbrite beads (Polysciences Inc., Warrington, PA, USA) for size reference and alignment purposes. All flow cytometry files were analyzed using FlowJo version 10.6.1 (Flowjo LLC, BD Life Sciences, Ashland, OR, USA).

### DNA Sequencing

To look for genetic mutations between the dark-trained and parental *Prochlorococcus* NATL2A, genomic DNA was isolated for Illumina sequencing from biological duplicate cultures. Cells from 5 mL of culture were first pelleted by centrifugation at 12,000 ×*g* for 10 minutes. Genomic DNA was extracted using a previously published phenol/chloroform procedure (Wilson 2001) with the addition of a 100% chloroform extraction at the end to remove residual phenol contaminates. To minimize shearing, samples were shaken briefly by hand (no vortex use), centrifugation time and speed (3 min at 16,000 ×*g*) were minimized, and used wide bore tips. Libraries were constructed using the NextEra XT kit (Illumina) on an automated Tecan Freedom EVO robotics platform, starting from 1ng of input DNA. The resulting libraries were sequenced using an Illumina NextSeq 500, generating between 7.6 million – 10.7 million 150+150nt paired-end reads from each sample (Supplementary Table 1). All library construction and sequencing was carried out by the MIT BioMicro Center. Sequencing reads have been deposited to SRA under BioProject PRJNA669190 (Supplementary Table 1). Low-quality regions of sequencing data and Illumina adapter sequences were removed using Trimmomatic (V0.36) (Bolger et al. 2014). Illumina sequencing data was analyzed by the *breseq* algorithm (V0.32) in both the default and polymorphism modes (Deatherage and Barrick 2014), aligning the individual reads to reference genomes for *Prochlorococcus* NATL2A and *Alteromonas macleodii* MIT1002 (NCBI GenBank accession numbers CP000095.2 and JXRW01000000, respectively). >99% of reads mapped to the reference genomes in each library; *Prochlorococcus* NATL2A had 833-1074x coverage, while both the genome and plasmid of *Alteromonas macleodii* MIT1002 had similar coverage levels within each library (in the range of ~80-200x). The read mapping evidence for any potential differences noted in ‘consensus’ mode was manually examined across both biological replicates, and also compared to the parental strains. Potential polymorphisms in the population noted by *breseq* (which could be due to either actual biological differences or systematic sequencing error) were examined for any site with a difference in >10% of reads. 6mA and 4mC methylation patterns were determined using Pacific Biosciences sequencing of triplicate samples. Culture samples were prepared by pelleting 90 mL of exponentially growing cultures by centrifugation at 7,500 ×*g* for 15 minutes. The pellet was resuspended into 200 μl of pH 8 TE and stored at −80°C. DNA was extracted using the same method as previously described above. Library construction began with diluting 4μg of intact genomic DNA in 150 μl of TE buffer (10 mM TrisHCl, 1 mM EDTA, pH 8.5) and fragmenting to 10-12 kb lengths using a gTube (Covaris Cat# 520079). Fragmented DNA was concentrated by a 0.45X SPRI bead cleanup (Beckman-Coulter), eluted in 37 μl EB (10 mM TrisHCl pH 8.0) and processed using SMRTBellTM Template Prep kit 1.0 (PacBio Part# 100-259-100) to SMRTbell libraries, following manufacturer guidelines. Indexed blunt-end SMRTbell adapters (PacBio Part# 100-4666-000) were added for multiplexing. Multiple indexed SMRTbell libraries were assembled in a single pool, cleaned up using a Minelute reaction cleanup kit (QIAGEN Cat# 28204) to 100 μl volume in EB, followed by SPRI 0.4X cleanup, with final elution in 11 μl EB. Concentration and quality were assessed by Picogreen (ThermoFisher QuantIt Cat# P11496) and FEMTOpulse (Agilent). Pooled libraries were bound using Sequel Binding Kit 2.1 and sequenced for 10 h on the Pacific Biosciences Sequel I using a SMRTCell 1Mv2. Library construction and sequencing were carried out by the MIT BioMicro Center; detailed library statistics can be found in Supplemental Table 2. The Pacific Biosciences sequences have been deposited to SRA under BioProject PRJNA669190 (Supplementary Table 2). We looked for SNPs in the PacBio data as compared to the reference genomes based on mapping of consensus reads. Structural variant analysis was based upon comparisons using the median length subread for each polymerase read. Methylations in *Prochlorococcus* and *Alteromonas* Pacific Biosciences reads were identified in SMRT Link (V.6.0.0) using the base modification analysis with default parameters. This analysis was run with a reference file that contained both the *Prochlorococcus* NATL2A and *Alteromonas macleodii* MIT1002 genomes (as above). GFF files containing the methylations identified were parsed to create separate files for the methylations identified in *Prochlorococcus* and *Alteromonas.* MotifMaker (V0.3.1, https://github.com/PacificBiosciences/MotifMaker) was used to identify methylated motifs, if any, in the *Prochlorococcus* and *Alteromonas* methylation GFF files separately.

### Mass spectrometry-based methylation analysis

The detection and quantification of 6mA, 4mC and 5mC in genomic DNA was carried out using previously published methods (Boulias and Greer 2021). Cells from exponentially growing axenic biological triplicate cultures (7 mL of *Alteromonas* and 50 mL of *Prochlorococcus*) were pelleted by centrifugation at 7,412 ×*g* for 20-30 minutes. DNA from cell pellets and seawater media blanks was exacted using the Qiagen DNeasy Blood & Tissue DNA extraction kit (Cat# 69504). To free nucleosides, 0.5-1 μg of gDNA was digested using 10 U of DNA Degradase Plus (Zymo Research) in 30 μl reactions incubated for 2 h at 37 °C. After digestion the sample volume was brought to 100 μl with ddH20 followed by filtration using 0.22 μm Millex Syringe Filters (EMD Millipore). 5 μl of the filtered solution was analyzed by LC-MS/MS. The separation of nucleosides was performed using an Agilent 1290 UHPLC system with a C18 reversed-phase column (2.1 × 50 mm, 1.8 m). The mobile phase A was water with 0.1% (v/v) formic acid and mobile phase B was methanol with 0.1% (v/v) formic acid. Online mass spectrometry detection was performed using an Agilent 6470 triple quadrupole mass spectrometer in positive electrospray ionization mode. Quantification of each nucleoside was accomplished in dynamic multiple reaction monitoring (dMRM) mode by monitoring the transitions of 252.1 →136.0 for dA, 266.1→150.0 for 6mA, 228.2 → 112.1 for dC and 242.2 → 126.1 for 4mC and 5mC.

As a negative control in each UHPLC-ms/ms experiment, we included a “mock” digestion reaction, consisting of DNA Degradase Plus and digestion buffer in water, without any added DNA. This control established the background level of the nucleosides in the reagents, which was subtracted from the values obtained for each gDNA sample. The amounts of dA, 6mA, dC, 4mC and 5mC in the samples were quantified using corresponding calibration curves generated with pure standards and the ratios of 6mA/dA, 4mC/dC and 5mC/dC were calculated. Quantification of DNA methylation in samples required a measured value to be at least 2x above the mock background level.

## Results and discussion

### Marine picocyanobacteria can adapt to tolerate extended dark exposure

To see whether repeated periods of extended darkness enhances a cyanobacterium’s ability to recover from light energy deprivation, we subjected two strains of *Prochlorococcus* – MED4 (a HL-adapted strain) and NATL2A (a LL-adapted strain) – to total darkness for 3 days, brought them back into the light, allowed them to resume growth, and then repeated this cycle for multiple transfers (Fig. 1). Because our previous work demonstrated that *Prochlorococcus* could not survive more than 1.5 days of darkness in pure culture (Coe et al. 2016), these initial studies were carried out in co-cultures with the heterotroph *Alteromonas macleodii* MIT1002 (hereafter referred to as *Alteromonas*). After each exposure to darkness we measured the recovery time – i.e. the time it took for the culture to re-enter exponential growth as measured by bulk chlorophyll fluorescence (see Methods). Dark exposure was imposed in mid-exponential growth and transfers to fresh media were done when cells reached late-exponential phase. As is typical in these types of experiments (Coe et al. 2016), bulk chlorophyll fluorescence of the cultures decreased when cells were re-exposed to light, until the cultures resumed growth (Fig. 1).

**Figure 1.**
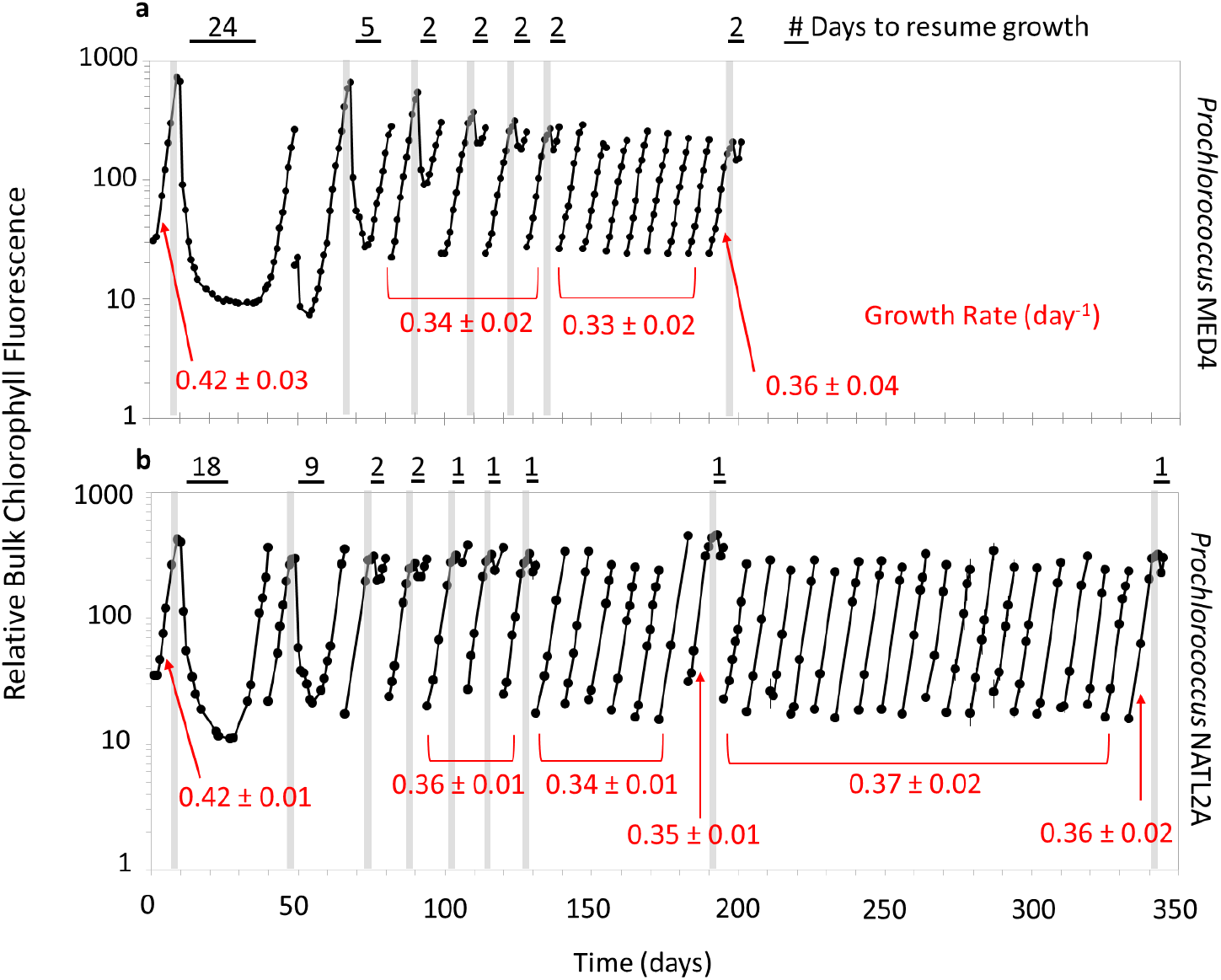
Recovery dynamics of *Prochlorococcus* cultures after repeated dark exposure. *Prochlorococcus* MED4 (a) and NATL2A (b) were subjected to 3 days of extended darkness (vertical gray bars) and then allowed to recover under standard 13:11 light:dark growth conditions. Once cells reached late-exponential growth phase, cultures were transferred to fresh media and the process was repeated. Transfers without vertical gray bars indicate growth under standard 13:11 light:dark conditions without extended darkness. The time needed to resume growth after dark exposure is indicated above the black horizontal bars and growth rates (day^−1^) calculated for the parental and dark-tolerant populations are shown in red. *Prochlorococcus* were grown in co-culture with *Alteromonas macleodii* MIT1002 to enable survival of the initial dark exposure.

Strikingly, after 3 rounds of repeated dark exposures and transfers, the time required to recover from dark exposure was reduced from 24 days to 2 days for MED4, and from 18 days to 1 day for NATL2A (Fig. 1). There were no further changes in the recovery time after the 4^th^ transfer. The MED4 cultures retained this ‘dark-tolerant’ phenotype for at least 7 transfers grown under standard 13:11 light:dark conditions (Fig. 1 a). NATL2A cultures were monitored for a longer period – an additional 18 transfers, or 33 generations, (Fig. 1 b, right) – under standard light:dark conditions. When then subjected to 3 days of darkness, these NATL2A cells still displayed the 1 day recovery time associated with the adapted, dark-tolerant phenotype. The shortened recovery times emerging from the ‘dark-training’ were not without a tradeoff: steady state growth rates (Fig. 1, red text) of the dark-tolerant cells during exponential growth were significantly lower than the parental ‘untrained’ culture (two-tailed *t* test, p < 0.05 for both MED4 and NATL2A). To explore whether this phenomenon might be a general feature of *Prochlorococcus*, we conducted a similar experiment with *Prochlorococcus* MIT9313 – a LL-adapted strain belonging to a different clade than NATL2A – and observed similar results (Supplemental Fig. 1).

To expand the taxonomic scope of the dark-tolerance phenomenon, we did a similar suite of experiments with marine *Synechococcus,* which has been shown to survive long periods of darkness in the wild (Sohrin et al. 2011). *Synechococcus* WH8102 can survive up to 7 days of darkness even without ‘helper-bacteria’ (Supplemental Fig. 2, red line), and can resume growth within 2 days when exposed to 3 days of darkness (Supplemental Fig. 2, black line). The latter is in striking contrast to *Prochlorococcus* strains which took 14-26 days to resume growth after their first exposure to 3 days of darkness (Fig. 1), indicating that *Synechococcus* has a different baseline tolerance of extended darkness. When we repeatedly exposed *Synechococcus* WH8102 to 6 days of darkness we observed the emergence of a dark-tolerant phenotype analogous to that seen in *Prochlorococcus*; dark recovery time decreased from 19 to 2 days after 3 cycles of darkness (Supplemental Fig. 3). The 2 day-recovery phenotype was stable through 7 transfers under standard 13:11 light:dark growth conditions, after which it was further reduced to 1 day (Supplemental Fig. 3, right side). While the growth rate of the *Synechococcus* dark-tolerant population was slightly lower than that of the parental cells, it was not significantly so (two-tailed *t* test, p = 0.2). Thus, while there some differences between *Prochlorococcus* and *Synechococcus vis a vis* the details of the dark-tolerance phenomenon, adaptation to repeated periods of extended darkness appears to be a general feature of picocyanobacteria.

### *Heterotroph interactions are not required for the dark-tolerance phenotype in* Prochlorococcus

As described above, the *Prochlorococcus* experiments could not be conducted with axenic cultures because the cells do not survive the first exposure to >1 day of extended darkness without the presence of a helper bacterium (Coe et al. 2016). That the axenic *Synechococcus* strain could exhibit a dark-tolerant phenotype suggested to us that heterotroph interactions might not be required for the phenotype itself, but instead were only important for extending dark survival long enough for the initial adaptation to occur. To test this hypothesis, we isolated lineages both the parental and dark-tolerant NATL2A cultures and did the same set of experiments with these axenic cultures. The axenic strain isolated from the dark-tolerant co-culture was still able to recovery rapidly from 3 days of extended darkness (Supplemental Fig. 4, black line), whereas the axenic strain isolated from the parental co-culture could not (Supplemental Fig. 4, pink line). Thus – as is the case for *Synechococcus –* the emergence of the dark-tolerant phenotype in *Prochlorococcus* does not require co-culture interactions. We corroborated this by isolating pure cultures of *Alteromonas* from the parental and dark-tolerant co-cultures and adding them back into axenic cultures of *Prochlorococcus* to see if they had any influence and there was no effect (Supplemental Fig. 4, red, blue, and gray lines). Thus dark tolerance emerges from changes within *Prochlorococcus*; the presence of *Alteromonas* is only needed to facilitate its survival during the initial selection period of 3 days of darkness.

### Dark-tolerance is associated with changes in population-level dynamics

In our previous work (Coe et al. 2016), we observed that when cells were put in extended darkness and re-exposed to light, chlorophyll fluorescence and light scatter per cell (a proxy for cell size) of each cell in the population decreased steadily over the recovery period until, eventually, a small population of about ~100 cells mL^-1^ (0.00015% of the cells that were placed in the dark) appeared with the same fluorescence and light scatter characteristics of the original population. We hypothesized that this represented the ‘seed population’ for the recovery of the culture, and that the remainder of the cells did not survive and/or recover. To probe this phenomenon in the dark-tolerant cultures, we examined flow cytometric signatures over the first 4 consecutive extended darkness ‘training’ cycles in *Prochlorococcus* MED4 (Fig. 2). During the first dark exposure and recovery (Fig. 2 a-f), the light scatter and fluorescence per cell decreased steadily over time and after 25 days (Fig. 2 e) the bulk of the population was near the baseline; a small population of cells however – here ~800 cells mL^−1^ – was evident with the fluorescence and light scatter values of the original parental cells (Fig 2 e, red dotted circle). This population appeared at the same time as the bulk chlorophyll fluorescence of the culture began to increase (Fig.1 a) – which is our metric for determining the beginning of recovery. With progressive cycles of dark exposure and recovery (transfers 2, 3, and 4, Fig. 2), the recovery time steadily decreased by halves, and a decreasing percentage of the population displayed the low fluorescence/light scatter stress response. By the 4^th^ cycle (Fig. 2 s-v), the recovery time had been reduced to 1 day (Fig. 2 u), the low light scatter/fluorescence population was nearly absent, and essentially the entire population retained the original ‘healthy’ flow cytometric signature. This indicates that after repeated ‘training cycles’ the vast majority of the population is able to maintain its chlorophyll and can ‘seed’ the recovery, explaining the decreased lag in regrowth: the seed population increased from 800 cells mL^−1^ (or 0.0006% of the cells that were placed in the dark) after the first round, to 4.5 × 10^6^, 9 × 10^7^, and 1.1 × 10^8^ cells mL^−1^ (or 6.19, 89.4, and 73.1% of the cells that were placed in the dark) in the 2^nd^, 3^rd^, and 4^th^ transfers respectively. The subpopulation with lower chlorophyll fluorescence likely reflects cells undergoing chlorosis, a degradation of the photosynthetic apparatus resulting in reduction of chlorophyll autofluorescence. Based on the regrowth patterns of the ‘seed’ population and observations that similar populations of 30-day old *Prochlorococcus* MIT9313 cells are inactive (Roth-Rosenberg et al. 2020), we hypothesize that the populations are likely non-viable. Thus, the subpopulation that is able to maintain a typical flow cytometric signature during extended darkness appears to be cells with the dark-tolerant phenotype.

**Figure 2.**
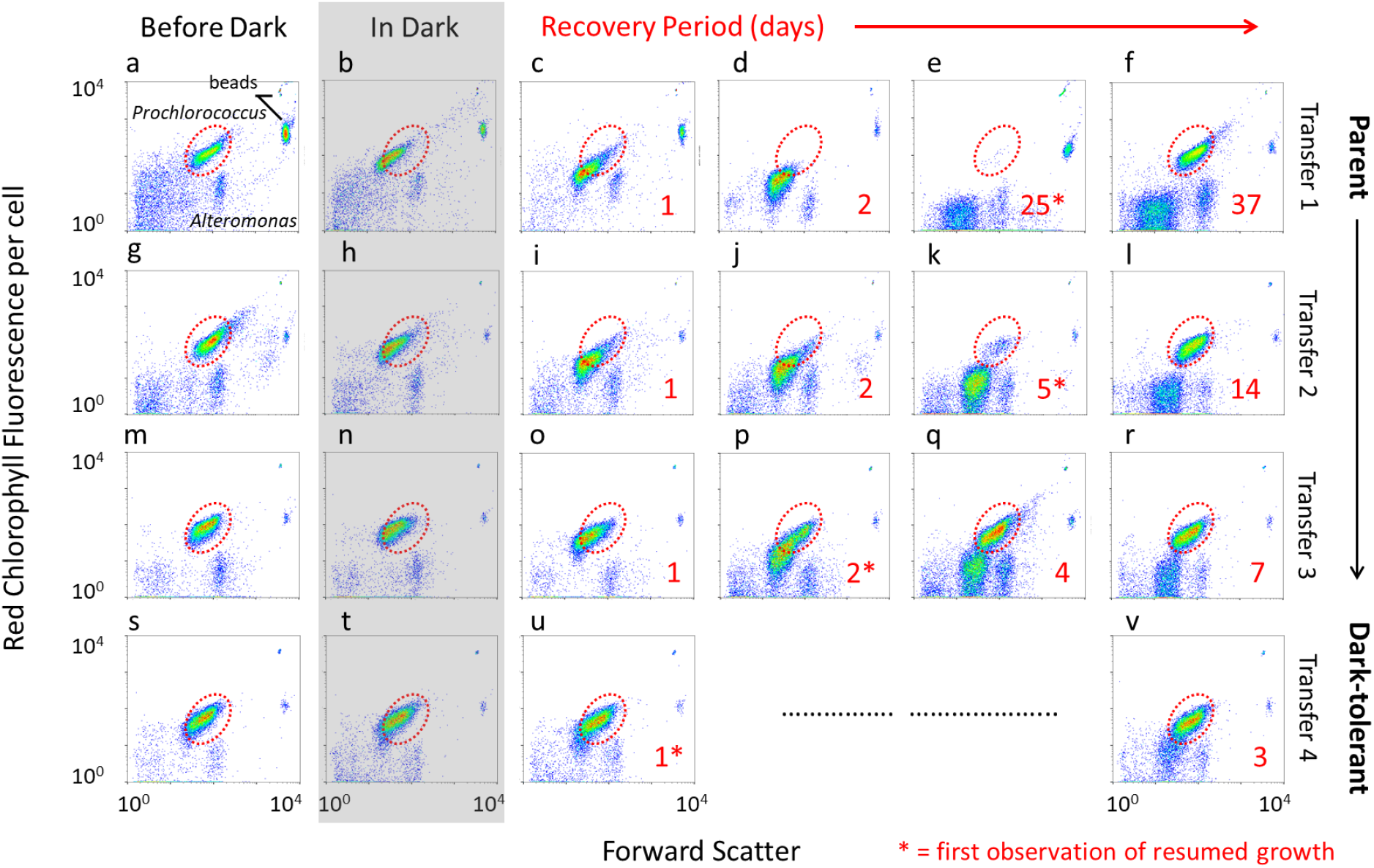
Flow cytometric analysis of *Prochlorococcus* MED4 populations during ‘dark-training’ by repeated dark exposure. Flow cytograms of forward light scatter and chlorophyll fluorescence per cell before dark exposure (first column), during 3 days of darkness (shaded area) and during the recovery period (panels to the right of shaded area) for each of 4 sequential transfers during the dark-training. Numbers in red denote the days during the recovery period (up to late exponential phase) and the first observations of resumed growth are indicated by (*) also seen as “days to resume growth” in Figure 1. The red dotted circle indicates the fluorescence and light scatter window occupied by parental and ‘fully recovered’ cells. The cultures were maintained under standard 13:11 light:dark conditions during the recovery periods.

### Dark-tolerant cells survive for longer periods of time in the dark

Dark-tolerant cells recover more quickly from extended darkness than do their parental cultures, but does this adaptation also confer an ability to survive longer periods of darkness? To examine this, we compared the recovery dynamics of wild-type *Prochlorococcus* MED4 cells (Coe et al. 2016) and dark-tolerant MED4 co-cultures to 3, 7, or 10 days of extended darkness (Fig. 3). MED4 parental cells could survive at least 7 days of darkness, but not more (Fig. 3). The dark-tolerant cells, however, were able to survive at least 10 days of darkness (longer periods were not tested). Furthermore, when the cells did recover, the speed of recovery was different between the parental and dark-tolerant cells (horizontal bars, Fig.3). Thus dark-tolerant *Prochlorococcus* cells likely can survive longer periods of time in dark and resume growth more quickly upon re-introduction to the euphotic zone than their ‘untrained’ counterparts.

**Figure 3.**
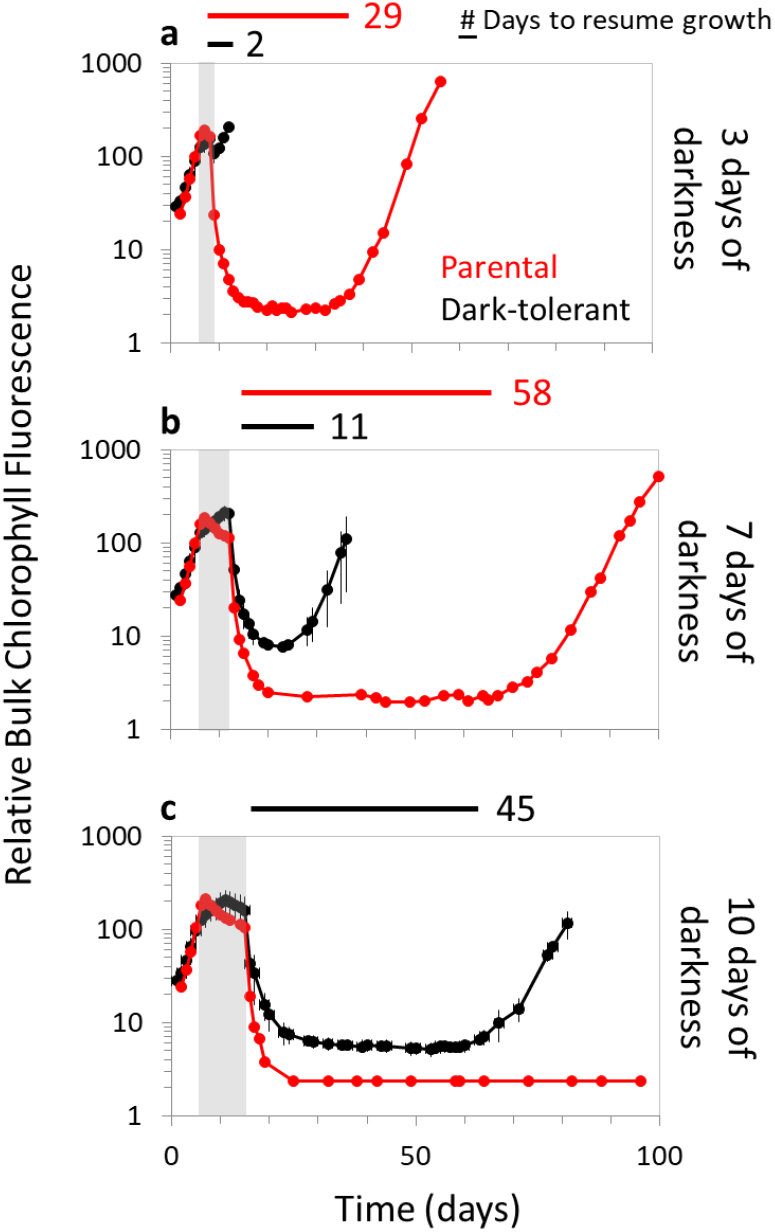
Response of dark-tolerant cells to increased durations of extended darkness. Parental (red) and dark-tolerant (black) cultures of *Prochlorococcus* MED4 grown in standard 13:11 light:dark conditions were subjected to either 3 (a), 7 (b), or 10 (c) days of extended darkness (vertical gray bars) and then returned to the light:dark cycle. The time needed to resume growth after dark exposure is indicated next to the horizontal bars. *Prochlorococcus* MED4 parental data (red lines) are from previously published data (Coe et al., 2016). *Prochlorococcus* were grown in co-culture with *Alteromonas macleodii* MIT1002 to allow survival of the dark exposure treatment.

### *Dark-tolerance is not due to a genetic mutation in* Prochlorococcus *NATL2A*

That the dark-tolerant phenotype was stable and heritable suggested that we had possibly selected for a genetic mutation within *Prochlorococcus*. To explore this we compared the genomes – including point mutations, insertions/deletions, gene duplications, inversions, or structural rearrangements – of replicate dark-tolerant *Prochlorococcus* NATL2A strains (sampled ~27 generations after the initial dark exposure) with those of the parental cultures using Illumina sequencing (~950x genome coverage; Supplementary Table 1). There was no evidence of genetic changes. There were also no significant changes in polymorphism frequencies within the dark-tolerant cultures compared to the parental cultures, or differences among the small fraction of reads (<1%) which did not align to the reference genomes. Given our sequencing depth, an undetected mutation in the dark-tolerant cultures would have to be present in less than ~0.1% of cells, which is not consistent with the observed bulk culture regrowth dynamics (Figure 1b). Additionally, long-read sequencing data obtained using Pacific Biosciences technology also failed to reveal evidence for large-scale chromosomal rearrangements or insertions/deletions in the dark-tolerant cells.

### *Potential for epigenetic regulation in* Prochlorococcus *NATL2A*

Given the lack of evidence for genetic change, we propose that the dark-tolerant phenotype in *Prochlorococcus* NATL2A must be mediated by an epigenetic mechanism. To explore this, we examined whether the dark-tolerance phenotype is associated with changes in DNA methylation. Many microbes contain DNA methyltransferases which add methyl groups to specific positions on DNA (Blow et al. 2016). These chemical modifications can disrupt DNA-protein interactions, serving to influence phage defense through restriction-modification systems (Tock and Dryden 2005), but can also alter transcriptional regulation (Mouammine and Collier 2018; Casselli et al. 2018). Indeed, DNA methylation-based changes in gene expression have been observed in the response of the freshwater cyanobacterium *Synechocystis* to nitrogen starvation (Hu et al. 2018), and the strains in our study encode at least one putative DNA methyltransferase enzyme (Roberts et al. 2010; Malmstrom et al. 2010; Stucken et al. 2013). Thus we examined the Pacific Biosciences sequencing data we had in hand from the genome sequencing efforts to look for DNA modifications in either the parental or dark-tolerant *Prochlorococcus* NATL2A co-cultures (i.e. Fig. 1 b, after both the 1^st^ and 7^th^ transfers). The single-molecule sequencing data revealed no methylated sequence motifs in either the parental or dark-tolerant *Prochlorococcus* NATL2A; as an internal control, we note that methylated motifs indicative of an active Dam methylase was found in the co-cultured *Alteromonas* genome.

To determine whether *Prochlorococcus* could have used a DNA modification that was not detectable by Pacific Biosciences sequencing, we extended our sequencing-based analysis with a mass spectrometry approach (O’Brown et al. 2019) to quantify the abundance of methylated nucleotides in our strains. Again, methylated nucleotides were not detected in either the parental or dark tolerant strains, while – again as a positive control – N^6^-methyladenine (6mA) methylations were detected in *Alteromonas* (Table 1). Surprised by the absence of methylations in this particular strain of *Prochlorococcus,* we wondered if any *Prochlorococcus* strains methylate their DNA thus we examined at a suite of strains representing different HL and LL adapted *Prochlorococcus* clades (Table 1). Both N^6^-methyladenine (6mA) and C^5^-methylcytosine (5mC) modifications are evident (Table 1) and differences in the type of methylated nucleotides can be seen across the strains. Differences in the relative fraction of methylated nucleotides between members of the HL and LL clades can be seen as well (Table 1). Thus, while select *Prochlorococcus* are able to methylate their DNA, this does not appear to be the causal mechanism underlying the dark-tolerance adaptation in NATL2A strain used in our experiments as it showed no evidence of methylation.

**Table 1.** Relative abundance of methylated DNA nucleotides in different *Prochlorococcus* strains, as well as *Alteromonas,* determined by mass spectrometry. N^6^-methyladenine (6mA), C^5^-methylcytosine (5mC), and N^4^-methylcytosine (4mC) methylated nucleotides were measured in axenic parental (white) *Alteromonas* and *Prochlorococcus* strains from different high-light (HL) and low-light (LL) clades grown under standard light:dark conditions and in axenic dark-tolerant (gray) *Prochlorococcus* NATL2A after experiencing changes in light regimes (treatments) including: before dark, 3 days in darkness (‘in dark’), and after the cells resumed growth and reached late-exponential growth phase (‘recovered’). Nucleotides that were not detected were indicated with ‘n.d’ and values indicate the mean (± sd) from three biological replicates.

| Strain | Treatment | Clade | 6mA/dA (%) | 5mC/dC (%) | 4mC/dC (%) |
| --- | --- | --- | --- | --- | --- |
| *Prochlorococcus* |  |  |  |  |  |
| NATL2A | - | LLI | n.d. | n.d. | n.d. |
| Dark-tolerant NATL2A | Before dark | LLI | n.d. | n.d. | n.d. |
| Dark-tolerant NATL2A | In dark | LLI | n.d. | n.d. | n.d. |
| Dark-tolerant NATL2A | Recovered | LLI | n.d. | n.d. | n.d. |
| MED4 | - | HLI | 1.605 (±0.08) | n.d. | n.d. |
| MIT9312 | - | HLII | n.d. | 0.847 (±0.029) | n.d. |
| MIT9215 | - | HLII | n.d. | 0.895 (±0.019) | n.d. |
| MIT9202 | - | HLII | n.d. | 0.318 (±0.017) | n.d. |
| AS9601 | - | HLII | n.d. | 1.006 (±0.044) | n.d. |
| MIT1304 | - | LLII/III | 0.317 (±0.007) | n.d. | n.d. |
| MIT9313 | - | LLIV | 0.063 (±0.02) | n.d. | n.d. |
| *Alteromonas* |  |  |  |  |  |
| MIT1002 | - | - | 7.708 (±0) | n.d. | n.d. |
| Dark-tolerant MIT1002 | Before dark | - | 8.455 (±0.003) | n.d. | n.d. |

DNA methylation is not the only possible heritable epigenetic modification that could be responsible for the phenotype. In fact, because *Prochlorococcus* contains relatively few protein-based transcription factors, mechanisms that do not typically work by impacting protein-DNA interactions might be more effective. Protein-based inheritance mechanisms in bacteria include, for example, systems with prion-like properties (Yuan and Hochschild 2017) and those based on the biased inheritance of protein abundance or modifications (Harvey et al. 2018). In addition, bacterial circadian clocks, first discovered in the cyanobacterium *Synechococcus elongatus*, can pass on the “time” to daughter cells via protein phosphorylation levels (Mihalcescu et al. 2004; Amdaoud et al. 2007). RNA modifications are yet another avenue for microbial epigenetics, as methylation of tRNAs has been implicated in bacterial translational regulation (Schwartz et al. 2018). Genetic feedback loops, in which initially small stochastic variation in gene expression from a ‘noisy’ promoter is ultimately amplified (or repressed) to influence genetic regulation, are also known to lead to heritable, epigenetic changes (Turner et al. 2009). These are all avenues for future investigations.

Finally, if the dark-tolerant phenotype we observed is indeed epigenetic, one might expect that the adaptation should be reversible on some observable timescale (Walworth et al. 2020). Indeed, following independent triplicate cultures of the NATL2A dark-tolerant phenotype under standard diel growth conditions for 195 generations, we found that 2 of the 3 cultures reverted after 55 months; they were no longer able to recover as quickly from extended dark exposure (Fig. 4, red lines) as they were after the initial training, however. Because this reversion occurred roughly simultaneously in two independent cell lineages, we suggest that epigenetic reversion is a more likely explanation than a compensatory genetic mutation, which might be expected to occur more stochastically. This reversion occurred only in the dark-tolerant co-cultures (Fig. 4, red lines) and not with the lineages that were rendered axenic (Fig. 4, blue lines), raising the possibility that some co-culture interactions facilitated the epigenetic switch back to the parental state – or that it simply has not yet occurred in the axenic triplicates.

**Figure 4.**
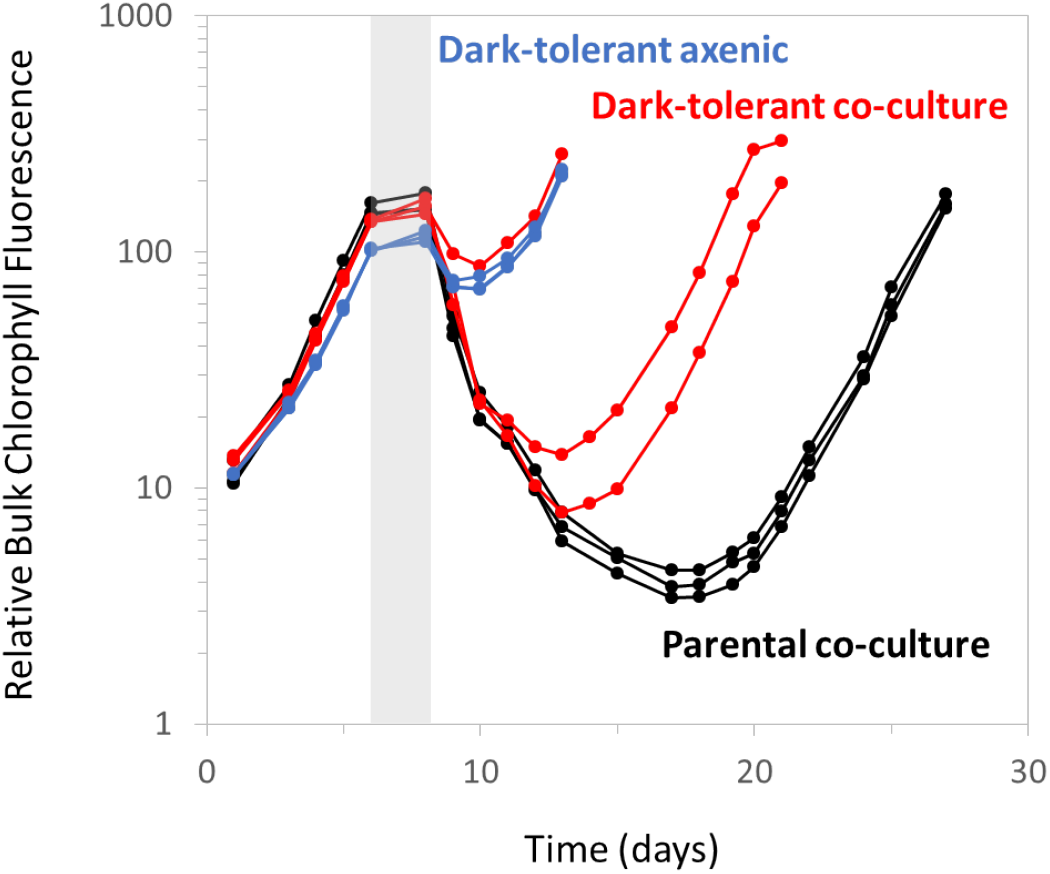
Spontaneous reversion of dark-tolerant *Prochlorococcus*. After 195 generations of growth under standard 13:11 light:dark conditions, the responses to dark exposure of individual triplicate lineages of dark-tolerant *Prochlorococcus* NATL2A co-cultures (red) were compared to those of the parental co-cultures (black) and axenic dark-tolerant cultures (blue). Cultures were subjected to 3 days of extended darkness (vertical gray bar), and their recovery under standard 13:11 light:dark conditions was monitored via bulk chlorophyll fluorescence.

## Conclusions

Subjecting a number of strains of *Prochlorococcus* and *Synechococcus* to repeated rounds of extended dark exposure selects for a heritable and reversible ‘dark-tolerant’ phenotype which evidence suggests is due to an as yet undescribed epigenetic mechanism. We propose that the ability to adapt to extended periods of darkness is likely a general feature of marine picocyanobacteria, adding a new dimension to the ecology of these globally abundant primary producers. A field study that resonates with our laboratory observations found that cells harvested from 397m in the Surgua Bay resumed growth within 2 days when exposed to light (Sohrin et al. 2011). This makes one wonder: Are most picoccyanobacteria in the oceans already ‘dark-tolerant’? Conversely, does maintaining cultured isolates under continuous light or diel light:dark cycling select for a ‘dark-sensitive’ state – the state of the parental strains in our experiments? At what frequency does the dark-tolerant phenotype occur in different habitats? Is it absent in areas of the oceans that are highly stratified throughout the year?

While our experiments revealed small fitness tradeoffs for *Prochlorococcus* in adapting to be dark-tolerant the potential benefit to the cell in regions where extended dark exposure is frequent is increased probability of survival to see another day. There may also be benefits accrued to the global *Prochlorococcus* ‘federation’ (*sensu* Biller et al. 2015) through this mechanism. Small sub-populations of dark-tolerant cells could be transported by deep currents to new locations, seeding surface populations with a new gene pool as a consequence of upwelling. As there is evidence for frequent exchange of genes among these populations (Biller et al. 2015), these ‘variants’ could introduce new genotypes for selection to act upon.

While our findings raise more questions than they answer, the dark-tolerant phenotype is quite striking and consistent – leading one to wonder whether similar adaptive responses to repeated stress stimuli await discovery, and whether epigenetic phenomena might play a more general role in shaping the ecology and evolution of marine microbes. The long generation times of bacteria from this habitat, coupled with the required length of these types of laboratory experiments means that progress will be slow. But it is well worth pursuing.

## Acknowledgements

The authors thank past and present the members of the Chisholm Lab for support and comments, especially Sean Kearney and Rogier Braakman for their valuable input. The authors would also like to thank Zev Cariani for assistance with sampling. We also appreciate sequencing efforts from the MIT BioMicro Center. This work was funded by grants from the Simons Foundation (Life Sciences Project Award ID 337262, S.W.C.; SCOPE Award ID 329108, S.W.C.). Work in the Greer lab was supported by a National Institutes of Health grant (DP2AG055947 to E.L.G.). We declare no conflicts of interest in performing and reporting these experiments.

