## Supplemental figures and tables for "Coping with darkness: The adaptive response of marine picocyanobacteria to repeated light energy deprivation"

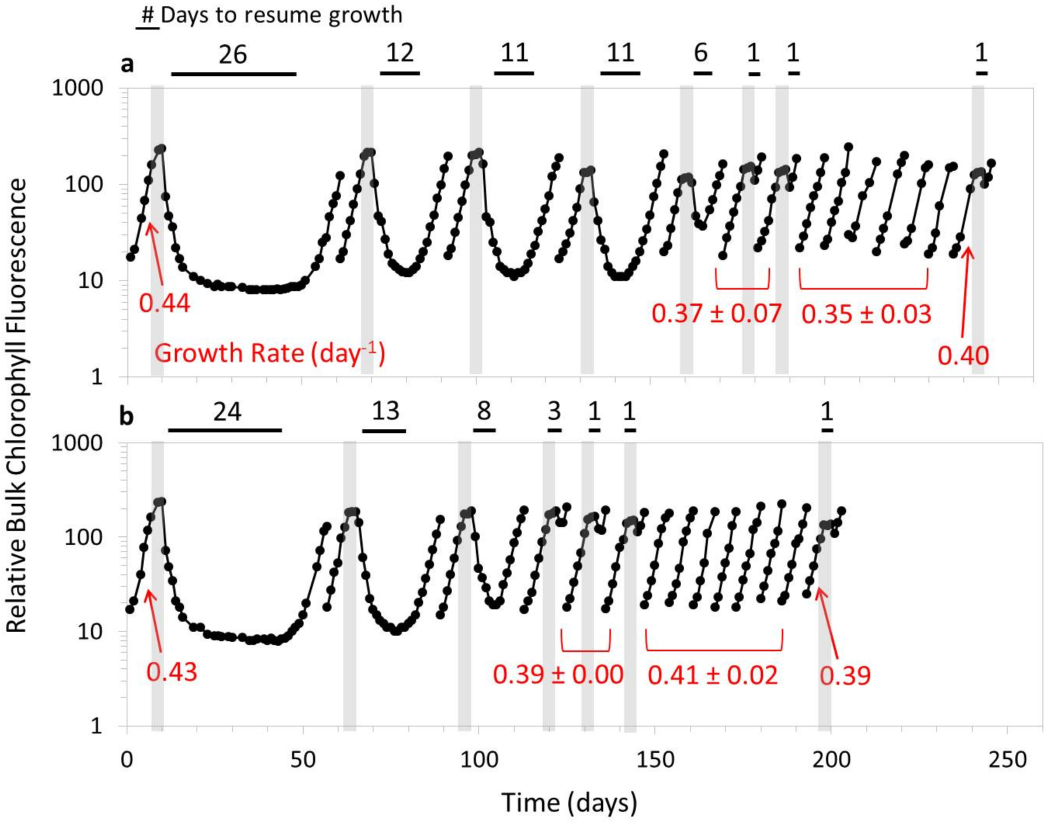


Supplemental Figure 1. **Response of *Prochlorococcus* MIT9313 to repeated light energy deprivation.** Single biological replicates (A & B) of *Prochlorococcus* MIT9313 were subjected to 3 days of extended darkness (vertical gray bars) and then allowed to recover under standard 13:11 light:dark growth conditions. Once cells reached late-exponential growth phase, cultures were transferred to fresh media and the process was repeated. Transfers without vertical gray bars indicate growth under standard 13:11 light:dark conditions without extended darkness. The number of days the cultures took to resume growth is above the black horizontal bars and growth rates ($day^{-1}$) calculated for the parental and dark-tolerant populations are shown in red. *Prochlorococcus* MIT9313 was co-cultured with *Alteromonas macleodii* MIT1002 to ensure survival of the first dark exposure.

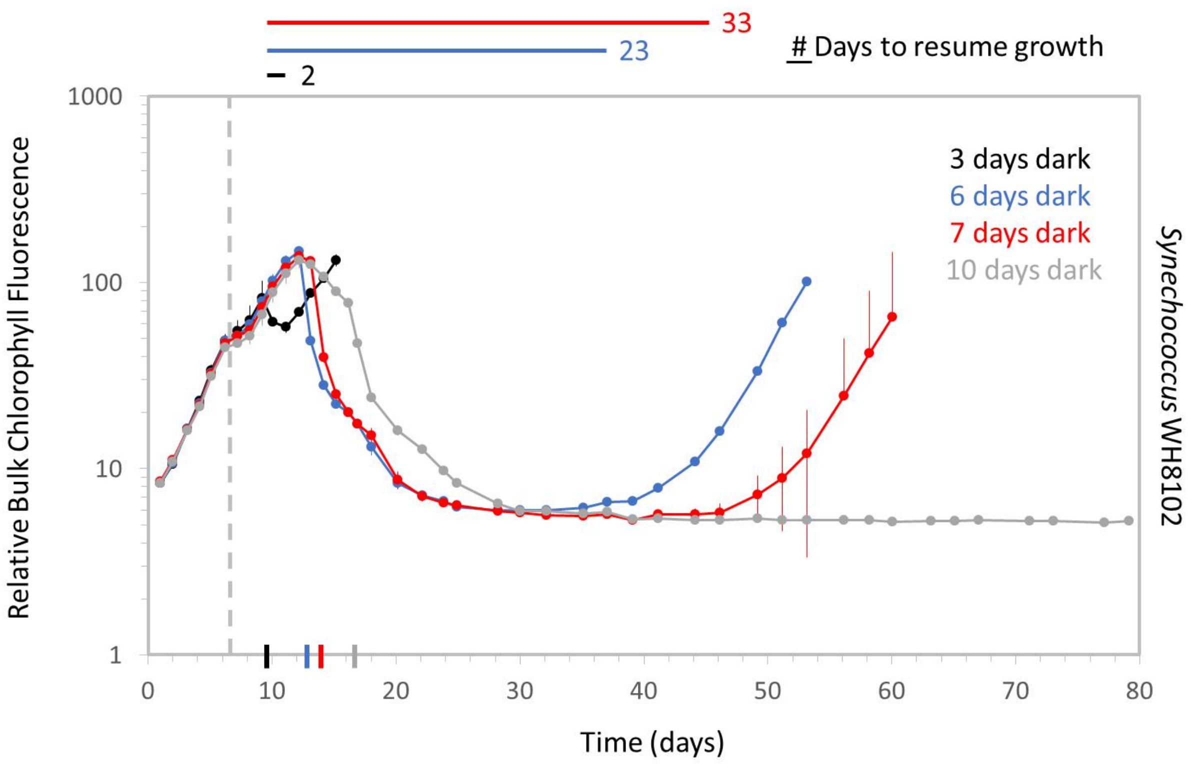


Supplemental Figure 2. **Response of axenic marine *Synechococcus* to extended darkness**. Replicate cultures grown on a standard 13:11 light:dark regime were grown to mid-exponential and then placed into darkness (gray dotted vertical line) for 3 (black), 6 (blue), 7 (red), and 10 (gray) days of darkness and then re-exposed to light:dark conditions (represented by colored ticks on axis). Cultures were monitored using bulk chlorophyll fluorescence for 80 days to detect if they could resume growth when placed back into the light (# days to resume growth next to horizontal bars); cultures which did not regrow during this timeframe were monitored visually for an additional 2 months, but no growth was observed.

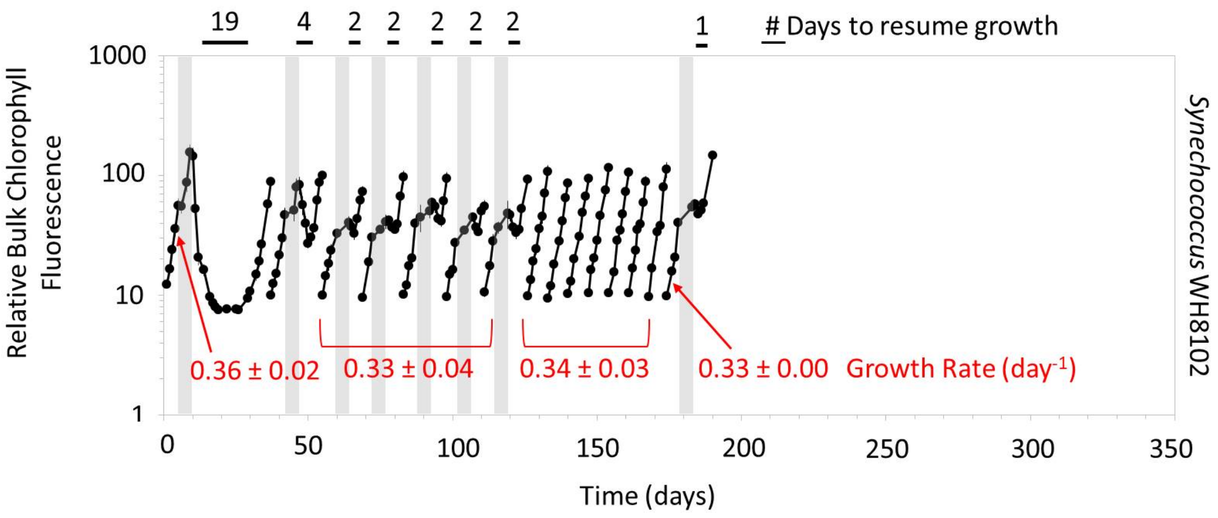


Supplemental Figure 3. **Response of *Synechococcus* to repeated light energy deprivation.** Biological replicates of axenic marine *Synechococcus* WH8102 were subjected to 6 days of extended darkness (vertical gray bars) and then allowed to recover under standard 13:11 light:dark growth conditions. Once cells reached late-exponential growth phase, cultures were transferred to fresh media and the process was repeated. Transfers without vertical gray bars indicate growth under standard 13:11 light:dark conditions without extended darkness. The number of days the cultures took to resume growth is above the horizontal black bars and growth rates (day$^{-1}$) calculated for the parental and dark-tolerant populations are shown in red.

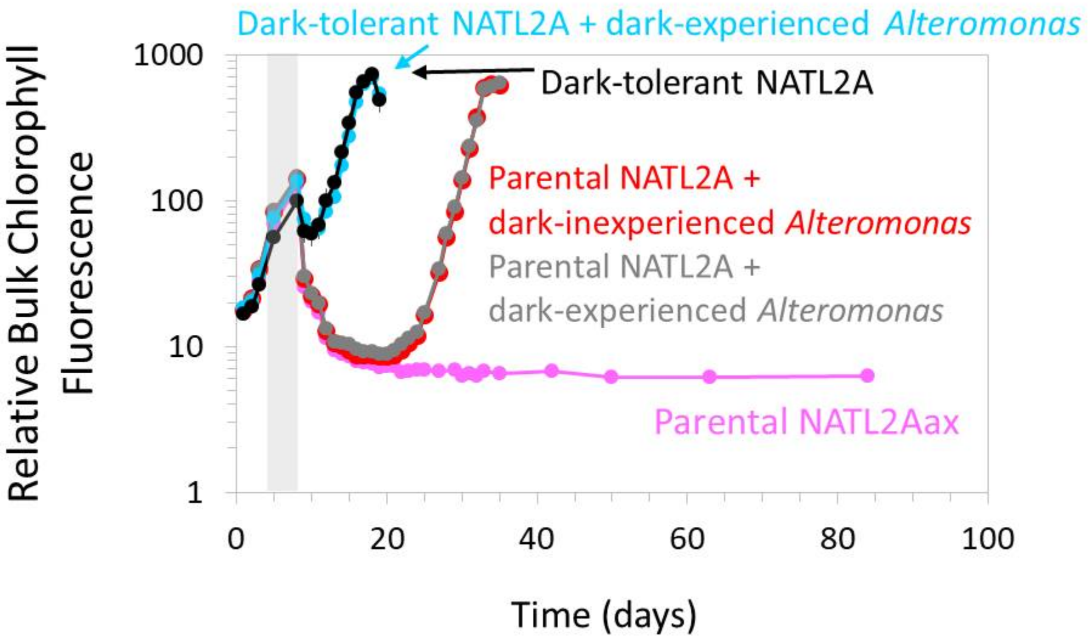


Supplemental Figure 4. **Role of *Alteromonas* in facilitating the recovery of *Prochlorococcus* from extended darkness.** Bulk chlorophyll fluorescence was monitored to follow the dark recovery response of axenic parental (pink) *Prochlorococcus* NATL2A when co-cultured with *Alteromonas macleodii* MIT1002 isolated from either dark-tolerant co-cultures (grey, 'dark-experienced') or parental co-cultures('dark-inexperienced, red). These responses were compared to a dark-tolerant culture recently rendered axenic (black) and dark-tolerant co-cultures (blue). All cells were placed in 3 days of darkness (vertical gray bar), after which cells were allowed to recover under standard 13:11 light:dark growth conditions.

Supplemental Table 1. **Illumina short-read sequencing datasets**

| Sample | Replicate | Total read pairs | Total bases | *Prochlorococcus* mean coverage | SRA Accession |
|---|---|---|---|---|---|
| Parent | 1 | 7,619,913 | 2,285,973,900 | 833x | PRJNA669190 |
| Parent | 2 | 10,740,362 | 3,222,108,600 | 1074x | PRJNA669190 |
| Dark-tolerant, after 1st transfer | 1 | 9,041,010 | 2,712,303,000 | 870x | PRJNA669190 |
| Dark-tolerant, after 1st transfer | 2 | 10,152,205 | 3,045,661,500 | 905x | PRJNA669190 |
| Dark-tolerant, after 7th transfer | 1 | 10,550,884 | 3,165,265,200 | 1028x | PRJNA669190 |
| Dark-tolerant, after 7th transfer | 2 | 10,296,848 | 3,089,054,400 | 1009x | PRJNA669190 |

Supplemental Table 2. **Pacific Biosciences long-read sequencing datasets**

| Culture | # Transfers / conditions | Polymerase Reads | Subreads | Total Size (bp) | Mean Read Length (bp) | Longest Subread Length (bp) | SRA Accession |
|---|---|---|---|---|---|---|---|
| Parent | 0 | 4,412 | 140,120 | 125,983,704 | 36,462 | 26,941 | PRJNA669190 |
| Parent | 0 | 14,219 | 662,706 | 469,641,058 | 38,889 | 28,868 | PRJNA669190 |
| Parent | 0 | 3,951 | 134,959 | 121,392,236 | 36,676 | 39,842 | PRJNA669190 |
| Parent | 7, normal light:dark cycle | 5,993 | 167,489 | 144,680,381 | 37,682 | 26,028 | PRJNA669190 |
| Parent | 7, normal light:dark cycle | 5,750 | 242,286 | 165,399,624 | 38,825 | 29,828 | PRJNA669190 |
| Parent | 7, normal light:dark cycle | 7,084 | 284,275 | 227,939,505 | 37,176 | 25,828 | PRJNA669190 |
| Dark-tolerant | 7, extended darkness | 19,117 | 1,000,335 | 651,250,400 | 40,707 | 16,276 | PRJNA669190 |
| Dark-tolerant | 7, extended darkness | 19,676 | 933,871 | 639,316,105 | 39,638 | 68,344 | PRJNA669190 |
| Dark-tolerant | 7, extended darkness | 20,722 | 1,089,743 | 697,667,938 | 40,301 | 42,026 | PRJNA669190 |
